# A multi-network approach to Alzheimer’s Disease gene prioritization complements *cis*-regulatory prioritization with molecular quantitative trait loci

**DOI:** 10.1101/2023.05.03.539189

**Authors:** Jeffrey L. Brabec, Montana Kay Lara, Anna L. Tyler, J. Matthew Mahoney

**Affiliations:** University of Vermont Department of Neurological Sciences 89 Beaumont Avenue Burlington, VT 05401; The Jackson Laboratory 600 Main Street Bar Harbor, ME 04609

## Abstract

Gene prioritization within mapped disease-risk loci from genome-wide association studies (GWAS) remains one of the central bioinformatic challenges of human genetics. This problem is abundantly clear in Alzheimer’s Disease (AD) which has several dozen risk loci, but no therapeutically effective drug target. Dominant strategies emphasize alignment between molecular quantitative trait loci (mQTLs) and disease risk loci, under the assumption that cis-regulatory drivers of gene expression or protein abundance mediate disease risk. However, mQTL data do not capture clinically relevant time points or they derive from bulk tissue. These limitations are particularly significant in complex diseases like AD where access to diseased tissue occurs only in end-stage disease, while genetically encoded risk events accumulate over a lifetime. Network-based functional predictions, where bioinformatic databases of gene interaction networks are used to learn disease-associated gene networks to prioritize genes, complement mQTL-based prioritization. The choice of input network, however, can have a profound impact on the output gene rankings, and the optimal tissue network may not be known *a priori*. Here, we develop a natural extension of the popular NetWAS approach to gene prioritization that allows us to combine information from multiple networks at once. We applied our multi-network (MNFP) approach to AD GWAS data to prioritize candidate genes and compared the results to baseline, single-network models. Finally, we applied the models to prioritize genes in recently mapped AD risk loci and compared our prioritizations to the state-of-the-art mQTL approach used to functionally prioritize genes within those loci. We observed a significant concordance between the top candidates prioritized by our MNFP method and those prioritized by the mQTL approach. Our results show that network-based functional predictions are a strong complement to mQTL-based approaches and are significant to the AD genetics community as they provide a strong functional rationale to mechanistically follow-up novel AD-risk candidates.

**Author Summary:** Risk genes give us insight into the failing molecular mechanisms that drive disease phenotypes. However, these risk genes are several layers of complexity removed from the emergent phenotypes they are influencing, the p-value that denotes their risk status gives little insight into the functional implications of that risk, and it is not clear *when* that risk gene may be having its effect. Methods like network-based functional prediction start to address several of these limitations by contextualizing risk genes in their broader genetic neighborhood within disease-relevant tissues. For complex diseases like Alzheimer’s, there are many possible relevant tissues incorporating everything from individual brain cell types to whole lobes of the brain. The work in this paper expands upon the traditional network-based functional prediction approach by considering a gene’s connections in multiple relevant tissue networks to prioritize candidate genes. Unlike traditional genetic risk studies, this kind prioritization benefits the Alzheimer’s genetics community as it provides a strong functional rationale to mechanistically follow-up on novel gene candidates.

## Introduction

Late-onset Alzheimer’s disease (LOAD) is a genetically complex neurodegenerative disorder characterized by progressive memory loss and cognitive dysfunction (1) involving all major cell types of brain (2). Dozens of genetic risk loci have been associated to LOAD-related disease processes through genome-wide association studies (GWAS) (3–9) and candidate genes within these loci have been nominated by multiple criteria (9–13). A subset of these genes has been mechanistically validated (14–16), but most LOAD risk loci only have putative candidate variants, and the functional role of these loci in LOAD pathogenesis remains unclear. Moreover, because GWASs use single-nucleotide polymorphisms (SNPs) that vary across the human population to identify disease risk loci by statistically associating SNPs to phenotypes, the causal effect of mapped SNPs on disease risk, if any exists, cannot be resolved from the GWAS data alone. The most statistically significant SNP in a region––the *index SNP*––may only be *linked* to the causal variant, meaning that it is only correlated with the presence of the truly causal variant or variants. Thus, the interpretation of SNP associations from GWAS requires *functional prioritization* to rank variants or genes near the index SNP as most likely to causally influence the phenotype. In general, the nomination of candidate genes within genetic risk loci is a complicated analytical process that requires a combination of auxiliary data integration, bioinformatic analysis, and biological prior knowledge (9,12,17–20).

One state-of-the-art strategy is to overlap genetic risk loci with expression quantitative trait loci (eQTLs), which are genetic associations to gene expression levels, presumably due to variation in *cis*-regulatory elements driving transcript-level variation. An eQTL-based candidate prioritization is predicated on a mechanistic model whereby a variant in a genomic sequence changes the expression level of a target gene, which then goes on to alter the risk for developing the disease. The eQTL-based approach generalizes naturally to other molecular modalities, including protein abundance QTLs, metabolomic QTLs, or epigenetic state QTLs. In the most up-to-date LOAD GWAS data from Bellenguez *et al.*, the authors performed a detailed QTL-based prioritization for 42 novel loci detected in their study, nominating 31 candidate genes (9). However, for the causal model underlying QTL-based prioritization to be valid, the molecular data set must match the correct cellular, tissue, and temporal context for the disease. Indeed, Connally *et al*. have recently performed a re-analysis GWAS data sets for seven complex traits that have cognate Mendelian forms with known gene mutations (21) and found “limited evidence that the baseline expression of trait-related genes explains GWAS associations”. Thus, they call into doubt the eQTL-based prioritization approach, which they term the *missing regulation problem*. One possible explanation for the missing regulation problem is that technical limitations, such as that molecular QTLs often derive from bulk tissue, obscuring a cell-type-specific context. Another possibility is that the altered regulation of the trait-relevant gene is a dynamic phenomenon, either developmentally or in response to an insult, but does not alter the baseline transcript abundance.

In this light, LOAD is a particularly challenging disease for eQTL-based prioritizations for two distinct reasons. First, symptoms emerge late in life after potentially decades of exposure to environmental risk factors and significant underlying tissue damage has accumulated (22). Second, genetic risk factors for LOAD act through complex neural circuits composed of many distinct cell types that interact dynamically over multiple time and length scales to encode memories and support cognition. Thus, for example, gene expression signatures in end-stage disease may not represent the mechanistically relevant time point. Moreover, while most of the brain is affected by LOAD, gene expression signatures might not measure the mechanistically relevant brain tissue. In many cases, this context is unknown and may even be inaccessible, such as during brain development.

A powerful complementary set of approaches to functional prioritization broadly form the category of *network-based functional prediction* (NBFP) (17,23–25). NBFP uses gene interaction networks to identify gene subnetworks that are enriched for disease-associated genes and to score all genes in the genome by their network connections to disease genes (17,23,24).

There are many gene interaction networks covering many contexts, including gene transcription networks (26), protein-protein interaction networks (27), and tissue-and cell-type-specific functional networks (28). For example, the popular NetWAS tool uses tissue-and cell-type-specific functional networks as input features for machine learning classifiers to rank genes according to their connectivity to disease risk genes. The choice of input network amounts to selecting a context in which causal variants are expected to be relevant, which can have a significant influence on the output gene rankings. For complex diseases, there may even be many relevant networks, e.g. multiple cell types of interest. Indeed, even the relevant cell types for LOAD remain controversial, and all cell types are likely involved in some potentially distinct capacity (29).

In this study, we extended the NetWAS approach (17), which only allows a single input network, to allow multiple input networks to aggregate evidence across many contexts to improve functional prioritization of GWAS candidate genes. We call our approach *multi-network functional prioritization* (MNFP). We applied MNFP with neuron and glia cell-type-specific networks from HumanBase 2.0 to LOAD-associated genes from the SNP-associations from the meta-analysis of Jansen *et al.* (8,28). Our MNFP model ranked all genes in the genome for mechanistic association to LOAD, and we compared these rankings to the corresponding single-cell-type rankings. We then ranked all genes in the newly identified LOAD GWAS loci from the Bellenguez *et al.* study and compared our top-ranking genes to corresponding QTL-based candidates (9).

## Results

### MNFP improves over single-networks to predict LOAD-risk genes

We performed four parallel NBFP runs using the three brain-cell networks, Neuron, Glia, and Astrocyte: one each for the dimension-reduced cell networks, and one for a multi-network fusion of those networks. During training, all network models performed approximately the same on average (astrocyte = 0.65, glia = 0.65, multi-network = 0.65, neuron = 0.62) with the neuron network having the lowest average AUC, which was significantly lower than the multi-network average AUC (*p* = 4e-8; Fig. 1A). In testing, the three single-network models performed, on average, the same as they did in training, with the neuron results showing modest improvement (astrocyte = 0.65, glia = 0.65, neuron = 0.64; Fig. 1B). In testing, the multi-network significantly outperformed each of the single networks, with an average AUC of 0.78 (neuron *p* = 1.2e-10, astrocyte *p* = 1.1e-8, glia *p* = 3.8e-9; Fig. 1B). Furthermore, across all outer CV folds, the multi-network had the lowest variance, 5.16e-4, of all the models (neuron = 5.89e-4, astrocyte = 8.6e-4, glia = 8.1e-4), suggesting that the multi-network produces more consistent models.

**Figure 1:**
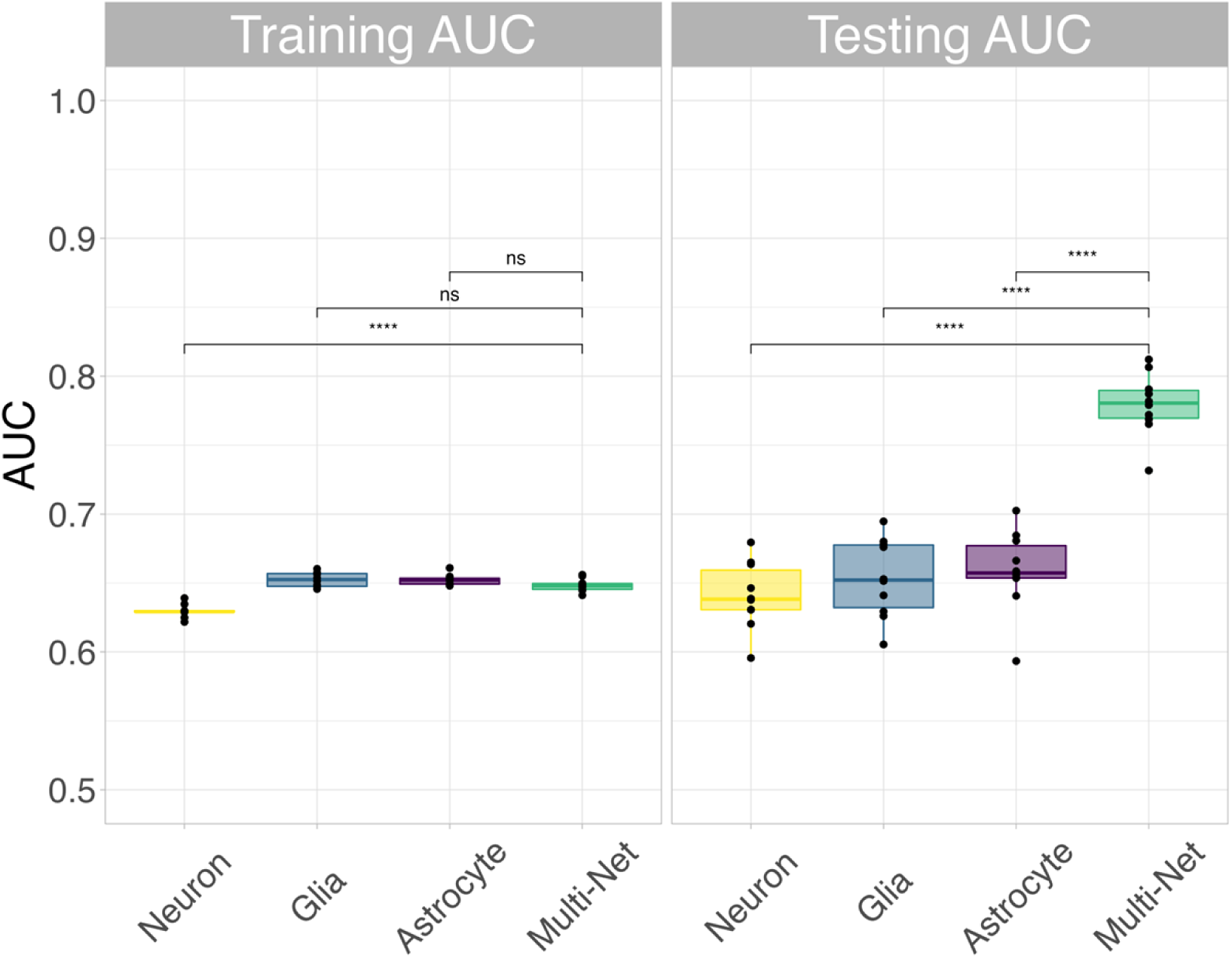
The NBFP model performance results. A) In training all models performed the same, on average (astrocyte = 0.65, glia = 0.65, multi-network = 0.65, neuron = 0.62). The MNFP performance was significantly better than the neuron performance in training (*p* = 4e-8). B) In testing, the MNFP significantly outperformed the single-network examples (neuron p = 1.2e-10, astrocyte p = 1.1e-8, glia p = 3.8e-9) the average performance of which stayed similar to the training performance (astrocyte = 0.65, glia = 0.65, neuron = 0.64). The MNFP approach also had the lowest performance variance, indicating more consistent models.

These results demonstrate that MNFP can more robustly predict LOAD risk gene networks compared to single-network NBFP. In particular, the increase in AUC for MNFP shows that the multi-network model has significantly fewer unlabeled genes (*i.e.*, non-true-positive genes) that are highly ranked by the SVM, suggesting increased confidence of the predicted disease association for those that are highly ranked compared to single-network models.

### Top Functional Candidates Are Network Hubs

To functionally annotate the pathways used by the multi-network model, we performed a modularity and enrichment analysis of the subnetworks of high-scoring genes for each source network. Spinglass-community detection and ontology analysis of the neuron network identified four modules of genes enriched for Gene Ontology terms involved in cell adhesion and motility, amyloid regulation, corticoid regulation, and immune regulation, respectively (Fig. 2). Each of these pathways are implicated in LOAD pathology, especially amyloid regulation. We sorted genes within each module by their degree to identify hub genes. The hub genes in the neuron network clusters were: *STK10* (cell adhesion and motility), *GRN* (amyloid regulation), *NRP2* (corticoid regulation), and *PHLDA1* (immune regulation). Interestingly, none of these genes were in the original true positive training set, though they appear to not only be hubs for their module but are densely connected to the other top functional candidates. The gene *NRP2* had the unique behavior of low functional scores in both the neuron (0.684) and glial (0.893) models and a good score in the astrocyte model (1.47), but a much larger score in the multi-network model (2.1). The additional information from all three networks has the potential to improve our candidate prioritization by nominating genes which may have important disease roles across several tissue types, but may be missed if only a single network is used where they play more minor roles.

**Figure 2:**
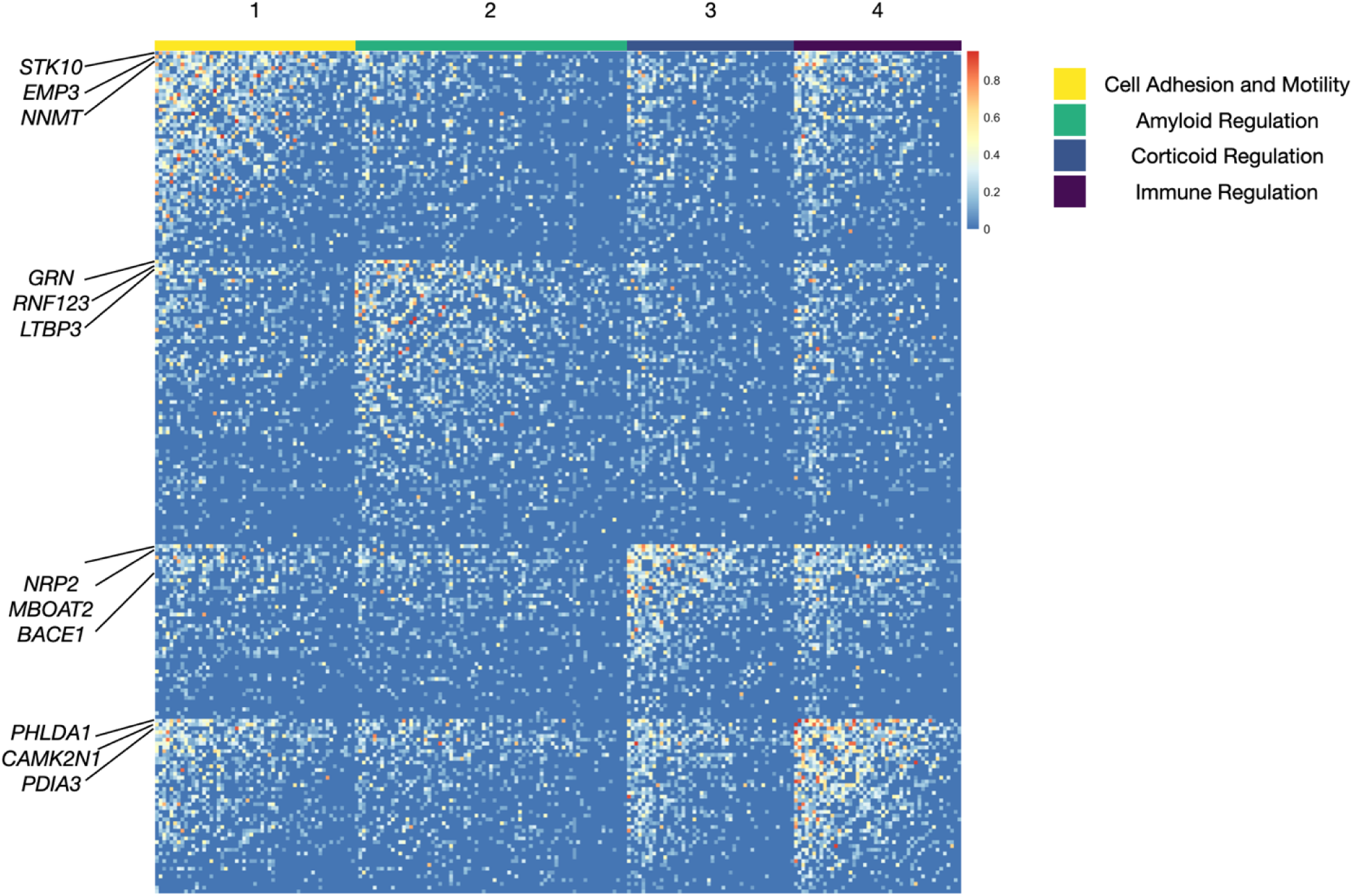
Heatmap of connections between top functional candidates in the neuron network. Spinglass-community detection identified four modules in the top neuron network. Genes in each module were ordered by degree. High-level summaries of the pathway enrichments of these modules indicate enrichments for cell adhesion and motility, amyloid regulation, corticoid regulation, and immune regulation. The top-degree genes of the cell adhesion and motility module include *STK10*, *EMP3*, and *NNMT*. The top amyloid regulation genes include *GRN*, *RNF123*, and *LTBP3*. Top corticoid regulation genes include *NRP2*, *MBOAT2*, and *BACE1*. The top-degree immune regulation genes are *PHLDA1*, *CAMK2N1*, and *PDIA3*.

Community detection and ontology analysis identified three modules in the astrocyte network involved in apoptosis, cellular responses to stress, and immune regulation, respectively. The hub genes for these clusters were: *IL12A* (apoptosis), *CCL2* (cellular response to stress), and *GREM1* (immune regulation) (Suppl. Fig. 1).

The same analysis identified three modules in the glia network involved in cell motility and adhesion, cell death regulation, and regulation of stress response, respectively. The hub genes for these clusters were *GREM1* (cell motility and adhesion), *IL12A* (cell death regulation), *DUSP5* (regulation of stress response) (Suppl. Fig. 2). Again, none of these hub genes were in the original set of true-positives used to train the model, though similarly to the top-astrocyte network, the hub genes *IL12A* and *GREM1* appear to be connected to a majority of the other top functional candidates in the glia network.

These results demonstrate that the network features used by multi-network NBFP to classify LOAD GWAS genes are enriched for LOAD-related biological processes and that the multi-network model assigns high scores to LOAD-relevant hub genes, including novel candidates that were not present in the training data.

### MNFP complements cis-regulatory predictions with molecular QTLs

Given recently reported associations of some non-true-positive hub genes to LOAD, we sought to systematically evaluate whether MNFP was able to identify novel LOAD gene associations. The recently published study from Bellenguez *et al.* implemented a novel functional prioritization method, briefly described in the methods section, to identify disease-gene candidates based on genome-wide significant risk variants (9). Their two-stage GWAS meta-analysis incorporated data from a number of different LOAD consortia (Methods). The Stage I analysis included 39,106 clinically diagnosed LOAD cases, 46,828 proxy-LOAD cases (based on a questionnaire asking whether the individual’s parents had dementia), and 401,577 controls. The Stage II analysis included 25,392 LOAD cases and 276,086 controls. All participants for this study were of European descent, or designated as non-white Hispanics in the incorporated datasets. They identified 42 genome-wide significant loci containing 55 novel putative LOAD gene associations which passed their prioritization criteria with 31 of those genes classified as Tier 1 and 24 as Tier 2. A gene received a Tier 1 ranking if the relative difference between its prioritization score and the other genes in its the locus was at least 20% and it obtained a score ≥ 4. If this threshold was not met, the highest ranked gene in the locus was assigned to Tier 2 (9).

To determine the concordance between our functional score and the candidates nominated by Bellenguez *et al.*, we scored all genes within the same 2 Mb windows to obtain an MNFP functional score-based ranking. The MNFP functional score is a continuous measure with no hard significance cutoff. Therefore, we assessed concordance of the two candidate prioritization methods by comparing the Bellenguez *et al.* top candidate(s) in each window to the top 1, 3, and 5 top functional score candidates (Table 1). To determine if the MNFP approach had greater concordance with the Bellenguez study than a single-network approach, we performed the same comparison with the single-network results. Comparison of a single top candidate per window revealed that our method shared 10 out of 55 top gene predictions with the Bellenguez *et al.* study (Table 2).

**Table 1:** Table summarizing concordant top-functional candidates between our method and that developed by Bellenguez *et al.* We examined overlaps for the top 1, 3, and 5 functional candidates.

| Network | 1 Top Candidate | 3 Top Candidates | 5 Top Candidates |
| --- | --- | --- | --- |
| Multi-Network | 10 | 17 | 22 |
| Neuron | 8 | 15 | 22 |
| Astrocyte | 8 | 19 | 24 |
| Glia | 4 | 15 | 23 |

**Table 2:** Summary table of concordant top functional candidates from our study and from Bellenguez et al.

| RSID | Interval | Our Top Candidate | Bellenguez *et al.* Top Candidate | Our Candidate's Functional Score | Bellenguez *et al.* Candidate's Functional Score |
| --- | --- | --- | --- | --- | --- |
| rs5848 | 17:43352876-45352876 | GRN | GRN | 2.06187503 | 2.06187503 |
| rs6586028 | 10:79494228-81494228 | TSPAN14 | TSPAN14 | 1.58058984 | 1.58058984 |
| rs2154481 | 21:25101558-27101558 | APP | APP | 1.46741222 | 1.46741222 |
| rs12592898 | 15:77936857-79936857 | CTSH | CTSH | 1.43481716 | 1.43481716 |
| rs7908662 | 10:121413396-123413396 | PLEKHA1 | PLEKHA1 | 1.314944 | 1.314944 |
| rs6584063 | 10:95266650-97266650 | BLNK | BLNK | 1.26932141 | 1.26932141 |
| rs1065712 | 8:10844613-12844613 | CTSB | CTSB | 1.26569387 | 1.26569387 |
| rs2245466 | 4:39197226-41197226 | RHOH | RHOH | 1.21893408 | 1.21893408 |
| rs1358782 | 20:0-1413334 | RBCK1 | RBCK1 | 1.03628506 | 1.03628506 |
| rs143080277 | 2:104749599-106749599 | NCK2 | NCK2 | 0.58929612 | 0.58929612 |

This concordance is highly non-random based on a permutation test where functional scores for genes in each locus are randomly permuted (*p* < 2e-5). (A permutation test was used because some windows overlapped, rendering analytic tests inapplicable.) Notably, for all 10 concordant candidates, the candidate gene was the closest gene to the index SNP. Beyond the top candidates, 17 candidates from Bellenguez *et al.* were ranked in the top three by multi-network functional score and 22 were ranked in the top five (Table 1). This high concordance with the Bellenguez *et al.* candidates was particularly pronounced in the multi-network model. Using the functional score from each of the single networks we obtained concordances of 8 (neuron), 8 (astrocyte), and 4 (glia) (Table 1). Together, these results demonstrate that the multi-network-based functional score is sensitive to functional information otherwise contained in direct data from patient brain tissue. Importantly, none of these top-5 candidate genes were part of the positive training set for the NBFP models, demonstrating that unlabeled-predicted positive genes from MNFP are highly enriched for functionally relevant LOAD genes.

In contrast to the concordant genes, there were 31 cases where the MNFP top-ranked gene had a functional score greater than one while the candidate from Bellenguez *et al.* had a functional score less than one (Table 3). Among these, several top candidates by multi-network NBFP have pre-existing literature evidence for association to LOAD such as *ARFGAP1*, *MBOAT2*, *MARCKS*, and *OAS1* (39–46).

**Table 3:** Divergent functional candidates between the two methods. Evidence for association to LOAD or another neurological disorder is provided by PubMed ID.

| RSID | Interval | Our Top Candidate | AD Associated Study (PMID) | Other Neuro/Metabolic Disorder (PMID) | Bellenguez *et al.* Top Gene | Our Top Gene's Functional Score | Bellenguez *et al.* Gene Functional Score |
| --- | --- | --- | --- | --- | --- | --- | --- |
| rs6742 | 20:62743088-64743088 | ARFGAP1 | 32872134, 29240824, 30886015 | 22423108, 34040545, 22363216 | LIME1 | 2.78 | 0.33 |
| rs9304690 | 19:48950060-50950060 | GYS1 | 22792253 |  | SIGLEC11 | 2.67 | 0.08 |
| rs149080927 | 19:854255-2854255 | GADD45B | 28050792, 28380666, 34842321 | 30832463, 25678854, 31043581 | KLF16 | 2.66 | 0.76 |
|  |  |  |  |  | REXO1 | 2.66 | 0.54 |
|  |  |  |  |  | ATP8B3 | 2.66 | 0.95 |
| rs35048651 | 17:728047-2728047 | PITPNA |  | 20471030, 16244667 | WDR81 | 2.54 | 0.96 |
|  |  |  |  |  | SERPINF2 | 2.54 | 0.76 |
| rs72777026 | 2:8558882-10558882 | MBOAT2 | 32109663 | 34742844, 22411505 | ADAM17 | 2.28 | 0.38 |
|  |  |  |  |  | ITGB1BP1 | 2.28 | 0.23 |
| rs2242595 | 17:17156140-19156140 | TOM1L2 | 34381170, 20167577, 34933910 | 32425817 | TOP3A | 2.18 | 0.62 |
| rs785129 | 6:113291731-115291731 | MARCKS | 34509657, 29557637, 33876503, 30155805, 10757536, 27557632, 31980612, 20651816, 11496150, 12865129, 25231903 | 30225354 | HS3ST5 | 1.90 | 0.04 |
| rs34173062 | 8:143103704-145103704 | OPLAH |  | 21653829 | SHARPIN | 1.86 | 0.52 |
| rs56407236 | 16:89103687-91103687 | TCF25 |  |  | PRDM7 | 1.84 | 0.22 |
| rs1140239 | 16:29010081-31010081 | SULT1A4 | 28374858 |  | DOC2A | 1.82 | 0.71 |
| rs587709 | 19:53267597-55267597 | PRKCG | 34254352, 32673674, 31441725, 25217572, 22369239, 17437536, 9576922, 11168531 | 30093405, 22369239 | LILRB2 | 1.80 | 0.48 |
| rs871269 | 5:150052827-152052827 | TCOF1 |  |  | TNIP1 | 1.79 | 0.78 |
| rs113706587 | 5:179201150-181201150 | CANX | 29725981, 34050864 | 34050864 | RASGEF1C | 1.67 | 0.20 |
| rs1160871 | 7:27129127-29129127 | HOXA5 | 33069246 |  | JAZF1 | 1.62 | 0.31 |
| rs16941239 | 16:85420604-87420604 | GSE1 |  |  | FOXF1 | 1.60 | 0.12 |
| rs6489896 | 12:112281983-114281983 | OAS1 | 34619763, 35383824, 35113409, |  | RITA1 | 1.46 | 0.70 |
|  |  |  |  |  | IOCD | 1.46 | 0.22 |
|  |  |  | 32274467,<br>33166007 |  |  |  |  |
| rs3822030 | 4:0-1993555 | MYL5 |  | 31766605,<br>29072034 | IDUA | 1.33 | 0.59 |
| rs17020490 | 2:36304796-38304796 | QPCT | 21900558 | 25848931,<br>26492838,<br>26572640,<br>34555062 | PRKD3 | 1.29 | 0.45 |
|  |  |  |  |  | CEBPZOS | 1.29 | 0.00 |
| rs3848143 | 15:63131307-65131307 | PPIB | 21683784,<br>21694447,<br>35305135,<br>25628064 | 27569281 | SNX1 | 1.28 | 0.39 |
|  |  |  |  |  | FAM96A | 1.28 | 0.00 |
| rs141749679 | 1:108345810-110345810 | GSTM2 | 35303681,<br>31628463 | 31628463 | SORT1 | 1.27 | 0.99 |
| rs112403360 | 5:13724304-15724304 | TRIO |  | 33183947,<br>32109419,<br>28928363,<br>26721934,<br>28973398,<br>31801062,<br>25363760,<br>25065758,<br>21422296,<br>25533962,<br>17043677 | OTULIN | 1.21 | 0.12 |
| rs7068231 | 10:59025170-61025170 | RHOBTB1 |  |  | ANK3 | 1.16 | 0.01 |
| rs76928645 | 7:53873635-55873635 | VOPP1 |  |  | EGFR | 1.02 | 0.14 |
| rs13237518 | 7:11229967-13229967 | THSD7A |  | 32627186 | TMEM106B | 1.01 | 0.84 |

## Discussion

In this study we developed MNFP as a simple and powerful extension of single-network techniques like NetWAS (17) for gene prioritization. Starting with a set of nominally significant GWAS genes, MNFP ranks all genes in the genome by how strongly they connect to these input genes in a functional network. This network-based ranking is distinct from the statistical signal that produced the input list (*e.g.*, GWAS *p*-value) and is often much more strongly enriched for disease-relevant biological signals (17,23). *A priori*, there are two criteria such a ranking should satisfy. First, the ranking should achieve good classification performance on the positive training inputs. This ensures that the prediction model has learned a biologically coherent signal from the potentially statistically noisy inputs. Second, the “false positive” predictions from the model should, in fact, be biologically meaningful inferences. In this study, we demonstrated that by combining information from multiple disease-relevant cell-type networks, we can improve over single-network NBFP on both criteria simultaneously; MNFP had a higher AUC than single-network models and was more highly concordant with functional prioritizations of gene candidates in novel LOAD risk loci than any single-network model (Table 1). By these benchmarks, our multi-network method is more conservative in predicting a LOAD association outside the input positive set, and such predictions more tightly align with functional prioritizations that account for extensive molecular evidence that is difficult and costly to generate (47). Not only did the multi-network approach perform significantly better than the single-network approaches, but it also had the lowest variability among all the models, showing that the multi-network is able to consistently build good models of functional significance. The success of MNFP in LOAD is a strong proof-of-concept for other complex genetic disorders.

In addition to methodological advances, the prioritizations of our multi-network model are of significant interest in LOAD. LOAD has become one of the most prevalent diseases of late life, but the genetic etiology is far from understood. Despite the significant concordance between our rankings with Bellenguez *et al.* in their recently identified risk loci, our method disagreed strongly with their candidate in 31 cases (Table 5). These discordant cases are potentially instructive about the strengths and weaknesses of the two approaches to candidate prioritization. The Bellenguez *et al.* approach emphasizes molecular QTLs that align with disease risk loci and *in vitro* gene knockdown screens of amyloid precursor protein (APP) metabolism as positive evidence of LOAD association. However, in LOAD, QTLs can only be ascertained in post-mortem brain tissue, which could miss a critical time point during development or pathogenesis when the causal gene is active. Likewise, a strong *in vitro* association with APP metabolism may be sufficient for functional relevance to LOAD but is not necessary. In contrast, our approach uses existing functional interaction networks that were trained using large compendia of mostly gene co-expression data (17,28). These networks effectively encode the emergent co-expression structure of different cell types and tissues, but do not have any *a priori* association to LOAD-specific biology. In ranking candidates in the Bellenguez *et al.* loci, our multi-network method seeks candidates within each locus that are strongly functionally connected to LOAD GWAS genes. Considering these differences, the overlap of ten genes as top candidates by both ranking approaches is highly surprising. Looking at these ten genes carefully, we see they have more in common than just top ranking in both methods: 1) each of these genes reached Tier 1 status (*i.e.*, strong evidence as a causal gene) by the Bellenguez *et al.* criteria, 2) each had a LOAD-associated eQTL, and 3) they all contained the index SNP defining their prioritization window. Interestingly, only one of the ten–– *APP––* had a measured effect on APP metabolism (9).

### Top Functional Candidates Mediating Amyloid Plaque Deposition

Of the ten genes share by both methods, *GRN* had the highest functional score and was a hub gene in the “amyloid regulation” module in the neuron network. The protein product of this gene, as well as its many smaller cleavage productions, are involved in a range of pathways including embryogenesis, tumorigenesis, inflammation, wound repair, and lysosome function (48). Variants in *GRN* are associated with a range of neurological disorders including fronto-temporal lobar degeneration (FTLD) and fronto-temporal dementia (FTD) (48,49). Loss of both alleles of this gene leads to the development of the disease neuronal ceroid lipofuscinosis (NCL), a lysosomal storage disease, which may contribute to LOAD pathology (48). It is hypothesized that high levels of Aβ can interrupt lysosomal signaling pathways, perhaps interfering with the mechanisms that would clear toxic protein aggregates (50). In a 2020 study, prior to *GRN* achieving genome-wide significance, researchers observed that *GRN* and several other sub-threshold LOAD genes showed increase expression in an APPswe/PS1^L166P^ mouse in response to high levels of Aβ (13). *GRN* is thus a strong functional candidate for LOAD.

The gene *APP* had the third highest functional score of the shared genes. *APP* codes for the protein which is eventually cleaved into Aβ and is therefore one of the most well-studied genes in LOAD (2,51–54). Several mutations in *APP* have been known to contribute to familial inheritance for years, but it only achieved genome-wide significance recently (6). Since *APP* is such a well-documented LOAD gene, we know that it is an integral part of disease subnetworks, with documented interactions with the biggest LOAD risk gene, *APOE* (55). *APP* is not a concordant gene in the neuron, glia, and astrocyte models until we compare the top 3 functional candidates. Thus, the strong MNFP score for *APP* demonstrates that integrating information from three different networks improves the identification of disease gene networks.

The other eight genes with shared top priority have been heterogeneously reported with respect to LOAD in the literature (Table 4). *CTSH*, *TSPAN14*, *NCK2* and *PLEKHA1* have all been implicated in LOAD or related disorders in recent genetic and protein screens (56–58). The genes *BLNK* and *CTSB* may act in the same pathway as *GRN* as they have both been observed to have altered expression in response to Aβ or roles in regulating lyosomal autophagy (13,59). Despite varying levels of literature evidence in support of roles in LOAD, the replication of these genes as top functional candidates in two independent studies warrants their further mechanistic study in LOAD pathophysiology.

### Divergence between MNFP candidates and cis-regulatory predictions

While a significant number of top candidates were shared between the two methods, there was a much larger set of functional predictions (n = 31) for which the two scores diverged (Table 5). To determine the biological differences between the two methods and the literature support for MNFP predictions, we performed a PubMed search in which we used the gene name with “Alzheimer’s” or “neuron” to identify genes which had previously been studied in the context of LOAD or played roles in other neurological disorders. Though there were many genes with neurological associations (Table 5), in the following we focus on those with the highest functional scores and strongest evidence for involvement in LOAD.

### MARCKS is an Early LOAD Gene

On chromosome 6, MNFP method prioritized *MARCKS* (FS = 1.90) in contrast to the Bellenguez *et al.* candidate HS3ST5 (FS = 0.04). *MARCKS* has been extensively studied in association with LOAD (Table 5) (41–43,60–63). *MARCKS* encodes the protein myristoylated alanine-rich C-kinase substrates that catalyzes cross-linking of actin filaments and is involved in diverse cellular pathways including cell adhesion and phagocytosis (62). Actin cross-linking is required for neurite outgrowth and dendritic spine stability. Phosphorylation of MARCKS is known marker of neurite degeneration in the early stages of Parkinson’s Disease (PD) and may be a viable marker for early-stage AD pathology (42,64). In LOAD, Fujita *et al.* have reported that in response to high levels of Aβ, glutamatergic neurons release the protein HMGB1, which binds to TLR4 and triggers the sustained phosphorylation of MARCKS causing neurite degeneration (43). They suggest that a monoclonal antibody targeting HMGB1 could be an effective treatment for preclinical AD to delay disease onset. In a subsequent study, the same group explored the broader effects of pMARCKS in cortical neurons of 5XFAD mice and observed decreased nuclear levels of Yes-associated protein (YAP) by immunohistochemistry due to increased sequestration in cytoplasmic amyloid plaques (63). YAP is a critical transcriptional regulator of homeostasis and its sequestration has been associated with early-AD necrosis (63). The Bellenguez *et* al. candidate in this locus, *HS3ST5* (heparan sulfate-glucosamine 3-sulfotransferase), has nine PubMed results, with only one of those containing evidence for a neurological association as a potential risk gene in a schizophrenia GWAS (65). The balance of evidence for this locus suggests that *MARCKS* is a superior functional candidate for the chromosome 6 locus, although we stress that absence of evidence is not evidence of absence and a significant mechanistic role for *HS3ST5* may emerge from further study.

### ARFGAP1 as a mediator of excitotoxicity in LOAD and PD

*ARFGAP1* (FS = 2.8) had the highest functional score of any Bellenguez *et al.* locus in our analysis, making it an excellent functional candidate for the chromosome 20 locus. The Bellenguez *et al.* candidate, *LIME1,* had a much low functional score (FS = 0.33). *ARFGAP1* codes for the protein ADP ribosylation factor GTPase activating protein 1 which is a Golgi apparatus-associated protein that hydrolyzes the ADP ribosylation factor 1-bound GTP (66). This activity is required for the proper dissociation of coat proteins, which help transport vesicles from the Golgi apparatus to the endoplasmic reticulum (66). *ARFGAP1* has been shown to be up-regulated in late stages of mild-cognitive impairment (MCI), though the precise functional implications of this finding are not clear (40).

*ARFGAP1*’s main neurological association is with PD. ArfGAP1 protein is known to physically bind to the kinase site of leucine-rich repeat kinase 2 (LRRK2) (67,68), whose kinase activity is a driving factor in the loss of dopaminergic neurons in PD (69,70). In 2012, Xiong *et al.* showed that LRRK2 and ArfgGAP1 have a reciprocal regulatory relationship. If expressed separately, ArfGaP1 or LRRK2 will induce cell death. However, if expressed together, ArfGAP1 binds to LRRK2 and promotes the hydrolysis of GTP which causes a decrease in LRRK2’s kinase activity and autophosphorylation. In turn, LRRK2 phosphorylates ArfGAP1 which inhibits further GTPase activation (67,68). Due to its high functional score and integral role in PD-related neurodegeneration, *ARFGAP1* is a strong risk gene candidate in the chromosome 20 locus.

### OAS1 Regulates Inflammatory Signals in Response to Aβ levels

The gene *OAS1* was the top functional candidate for the MNFP method (FS = 1.46) in the chromosome 12 locus. The corresponding Bellenguez *et al.* candidate was *RITA1* (FS = 0.70). *RITA1* has no literature association to LOAD or other neurological disorders. In contrast, Salih *et al.* recently nominated *OAS1* as a candidate risk gene for amyloid deposition based on increased expression of the mouse ortholog *Oas1a* in response to high amyloid levels (46). Lee *et al.* observed that *OAS1* has anti-inflammatory activity in macrophages, which is triggered through TLR3 and TLR4 signaling (44). It has been shown that microglial activation in LOAD is partly triggered through the binding of aggregated amyloid plaques to TLR4 receptors on microglia (71). Roy *et al.* showed that *Oas1*a is up-regulated in amyloid plaques and classified it as an “interferon stimulated gene” in their murine AD model (72). Magusali *et al.* showed that high levels of *OAS1* were critical for suppressing pro-inflammatory signals in myeloid cells (73). These results strongly implicate *OAS1* in LOAD-related neuroinflammation and support it as a functional candidate in the chromosome 12 locus.

### MNFP identifies disease genes across populations

As GWAS cohorts become larger, the power to detect risk variants increases, but the effect sizes for with these newly associated variants decreases. By design, MNFP aggregates over many weak-effect variants into a network model of pathogenesis. Beyond mapped risk loci, high-ranking genes from the MNFP model are also high-quality candidates for driving LOAD pathology. The fourth-ranked gene overall was the cytokine *IL12A* (FS = 3.70), which is a hub-gene in “apoptosis” and “regulation of cell death” modules in both the astrocyte and glia networks (Suppl. Fig 1 and 2). *IL12A* has primarily been studied for its role in cancers and auto-immune disorders (74–77), but it has recently also been associated with AD risk gene in a Han Chinese population (78). Notably, it has strong network connections to *OAS1*, discussed above, as well as *CXCL2*, another immune-modulating chemokine which is the second most densely connected gene in the astrocyte “apoptosis” module. Thus, our network results implicate *IL12A* as a potential immune effector of cell death in LOAD.

This result highlights an important distinction between a risk gene, which is at least partially dependent on the genetic background of a population, and a disease gene, which plays a mechanistic role in a disease. The goal of MNFP is to extrapolate from risk genes, given by GWAS associations, to a broader set of disease genes. Ideally, this extrapolation should lose the population specificity of its input and identify new genes that are nevertheless involved in the disease process. In the case of *IL12A*, MNFP extrapolated from risk genes derived from in a European-descent population to predict potentially novel disease genes, which, in this case, have polymorphisms influencing disease risk in the Han Chinese population.

### SPSB1 is a Candidate Mediator of MCI to AD Conversion

The overall top-scoring gene from our method was *SPSB1*, which codes for a protein that is involved in ubiquitination and promotes ubiquitin-dependent protein catabolism (79). Ubiquitination is a critical process for the clearance of misfolded or denatured proteins. As we age, the cellular systems that respond to misfolded proteins start to naturally decline and can be furthered impaired by pathogenic protein aggregates like amyloid and hyperphosphorylated tau (80). Two variants in *SPSB1* have been implicated in the conversion from MCI to LOAD (79). The conversion of MCI to AD is still poorly understood. Some patients develop MCI, but their cognitive dysfunction plateaus, while others have an ongoing degeneration leading to AD. The molecular mechanisms of this conversion are particularly important clinically, given that the onset of MCI is the earliest clinically relevant event for most patients with risk for developing AD. The strong functional association of *SPB1* to AD gene networks and the GWAS association to AD conversion implicates the misfolded protein response as a critical pathway for neurodegeneration specifically at this clinically actionable time point.

Taken together, our results show that by combining information across multiple cell-type-specific gene interaction networks, we achieve higher classification performance than single-network approaches. Our results implicate many strong gene candidates for LOAD pathology, both inside and outside genome-wide significant GWAS loci. These results complement *cis*-regulatory prioritizations, overcoming limitations due to sampling either healthy or end-stage-disease tissue, and strengthening the rationale for mechanistic studies of many novel genes.

## Methods

### Network Processing, Feature Selection, and Model Training

The network edge-lists for the neuron, glia, and astrocyte networks were obtained from the online repository HumanBase (https://hb.flatironinstitute.org/download; ‘neuron_top’, ‘glia_top’, and ‘astrocyte_top’). Briefly, these networks were generated using a regularized Bayesian knowledge integration based on tissue ontology and a combination of gene expression datasets from the Gene Expression Omnibus (30) representing 20,868 conditions (17). Each functional network is a weighted network, where each pair of genes (*g_i_*, *g_j_*) is linked with a weight, *W*_*gigj*_, encoding the predicted probability that those genes functionally interact in that tissue. We define a *feature vector*, *f_g_*, for each gene, *g*, in the genome as the vector of weights connecting *g* to the *n* LOAD-GWAS genes, *p*_1_,…,*p*_*n*_(*i.e.*, positive examples),

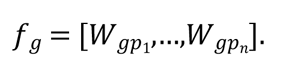

Each feature vector was dimension-reduced using a k-medoids method. In contrast to a k-means method, k-medoids chooses real data points as centers or medoids. Using this method, we reduced each feature set down to the 500 most representative genes. The dimension-reduced networks were then merged together to form the final multi-network feature set with 1500 features and 25,825 variables. In addition to creating the multi-network model, we processed each individual network identically into a dimension-reduced feature-set to generate single-network models. The performance of the single-network models, the more standard approach to NFP, was compared to the performance of the MNFP model to determine.

Using these feature vectors, we trained an ensemble of (linear) support vector machine (SVM) classifiers to distinguish between LOAD-GWAS (genes which had a GWAS *p <* 0.01) genes and the rest of the genes in the genome. Formally, this problem is an instance of *positive-unlabeled (PU) learning* (PU), as we only have positive examples of LOAD-relevant genes (*i.e.*, GWAS hits), but the status of all other genes is unknown. In the PU learning setting, we can treat all unlabeled examples as negatives for the sake of training the model, with the understanding that many unlabeled examples are likely LOAD-associated genes (31). We performed 10-fold nested cross-validation (CV) to ensure independent training and testing sets, as our positive examples our limited to the LOAD-GWAS genes. Hyperparameters were tuned in the inner CV loop and tested in the outer CV loop. The data were initially split into 10 equal folds, the outer loop, and stratified so that a proportional set of positive and unlabeled genes were present in each fold. Subsequently, each fold was split into testing and training sets. The training set of each fold was further split into 25 bootstraps, the inner-loop, again stratified on the class of each gene, which were also split into testing and training sets. The models in the inner-loop, were trained using a balanced set of positive and unlabeled examples (down-sampling the unlabeled set) as putative negatives. Each inner-loop model was cross-validated over the 25 bootstraps to optimize its cost hyperparameter, C, over a grid, as described previously (32). Each model *M_i_* assigns each gene, *g_j_*, a model-based, real-valued prediction score *M_i_*(*g_j_*), where scores above 0.5 correspond to high confidence that the gene is a positive example and scores below 0.5 correspond to low confidence. To determine a gene’s predicted score across all models, we calculated the logarithmic mean of its prediction score. To normalize prediction scores across models prior to aggregation, we computed an *unlabeled-predicted-positive rate* (UPPR) for each model, *M_i_*, and gene, *g_j_*, as

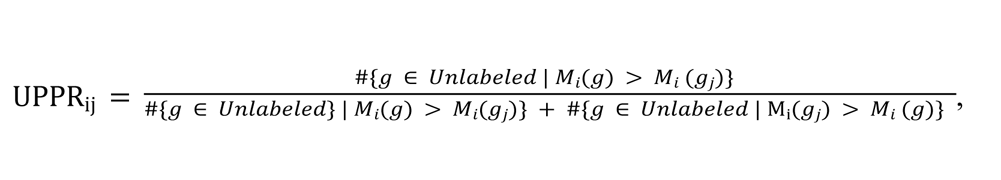

where ‘#’ denotes the cardinality of a finite set. The UPPR is the PU-learning equivalent of the false positive rate, where lower values indicate higher confidence that a gene is functionally associated with the LOAD GWAS genes. We took the average of UPPR over all models and took the negative logarithm to obtain a final *functional score*, *FS*(*g_j_*)

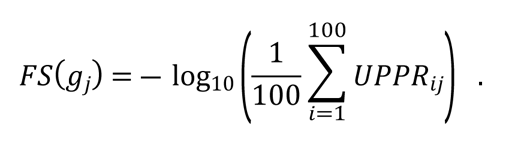

The functional score ranges from zero to infinity, with higher values indicating greater confidence. Models were trained using the *e1071* R package (33).

### Top Functional Network Analysis

NBFP identifies functional candidates based on their connections to the LOAD-GWAS genes. To determine how the top functional candidates interact with one another in the component cell-type networks, we performed module detection and pathway enrichment analysis. For all three cell-type networks, we extracted the subnetworks of genes which reached a functional score greater than two (*i.e.*, average UPPR < 0.01). Modules were identified using spinglass community detection implemented in R. The resulting networks were visualized using *pheatmap* and genes in each cluster were ordered by their degree (34). Pathway enrichment for Gene Ontology Biological Process (GO:BP) terms (35,36) for each module was performed using the g:GOSt function from the *gprofiler2* R package (37). High-level annotations for each cluster were annotated on the heatmap, as well as the top three genes in each cluster by degree.

### Functional Prediction Validation

The set of gold standard LOAD genes is dominantly GWAS risk genes, which makes it difficult to validate functional predictions. A recent LOAD meta-GWAS study performed functional prioritization using a novel method, and identified several novel LOAD candidate genes (9). Therefore, we decided to do a comparative analysis of the top functional candidates from our two studies. They performed a two-stage analysis which incorporated sample information from several consortia and datasets including: EADB, GR@ACE, EADI, GERAD/PERADES, DemGEne, Bonn, the Rotterdam study, the CCHS study, NxC, and UKBB for the Stage I analysis and ADGC, CHARGE, and FinnGenn for the Stage II analysis (9). Individuals for this study were of European descent, or designated as non-white Hispanics in the incorporated datasets. The Stage I analysis included 39,106 clinically diagnosed LOAD cases, 46,828 proxy-LOAD cases (based on a questionnaire asking whether the individual’s parents had dementia), and 401,577. The Stage II analysis included 25,392 LOAD cases and 276,086 controls.

In Stage I, a meta-analysis of the data from the included studies was conducted. They identified variants which reached a Stage I *p* < 10e-5, classifying them as an index variant, and defined a ±500 kb window around each. After merging the windows together, they performed an iterative clumping procedure to assign variants to clumps of the index variants, starting with the index variant with the lowest *p*-value. Variant which reached a Stage I *p* < 10e-5 and were located within the 1Mb window of the current index variant were assigned to that window. This process was repeated until all variants were assigned to clumps. For the Stage II analysis, results from the studies from both stages were combined in a fixed-effect meta-analysis. Follow-up variants needed to have the same direction of effect in both stages and a Stage II *p*-value < 0.05. Genome-wide significant candidates were identified from this set (n = 42 significant loci). Variant-to-gene mapping was performed using the MAGMA tool (38). Functional scores were calculated for genes within a 2Mb window of each significant locus. Their functional score incorporated: 1) variant annotation, 2) eQTL-GWAS integration, 3) splicing QTL (sQTL)-GWAS integration, 4) protein QTL (pQTL)-GWAS integration, 5) methylation QTL (mQTL)-GWAS integration, 6) histone acetylation QTL (haQTL)-GWAS integration, and 7) APP metabolism (9). Full descriptions of each domain can be found in their methods section.

Functional prediction resulted in 55 genes. We obtained summary statistics for these genes from their paper and replicated their 2MB window search, but instead prioritized by our functional score opposed to their QTL based method.

To compute a null distribution for the number of concordant top hits between the MNFP rankings and the Bellenguez *et al*. candidates, we performed a permutation test.

We randomly shuffled the functional scores of all positional candidate genes and calculated the number of shared hits. Because 100,000 permutations were insufficient to identify an overlap as large as observed––only 3% of all permutations had any overlap at all and no permutation exceeded 2 overlapping genes––we fit a Poisson distribution to the empirical null, from which we could calculate analytical p-values for any overlap size. Specifically, given the mean of the null distribution, we computed the analytic p-value of an overlap size.

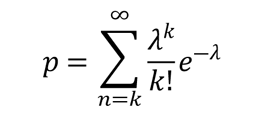

This formula approximates the probability of observing at least overlapping genes under the null hypothesis that the functional scores are random with respect to the Bellenguez *et al*. candidates.

## Data and Code Availability

The nominally significant LOAD GWAS genes obtained from the Jansen *et al.* study used to train the models can be downloaded from this link: https://ctg.cncr.nl/documents/p1651/AD_sumstats_Jansenetal_2019sept.txt.gz or found on their website https://ctg.cncr.nl/software/summary_statistics/. Summary statistics from the Bellenguez *et al.* paper were obtained directly from their paper. The code used to train the models and perform the analyses discussed in this section can be found here: https://github.com/jeffreyLbrabec/multi_network_prediction.

## Acknowledgements

The authors would like to acknowledge Rod Scott M.D./Ph.D., Amanda Hernan Ph.D., James Stafford Ph.D., Dimitry Krementsov Ph.D., and Josh Bongard Ph.D. for their guidance that helped to shape the narrative of this paper.

## Notes

### Competing Interest Statement

The authors have declared no competing interest.

